# Physics-informed stereology for estimating placental diffusive exchange capacity

**DOI:** 10.64898/2026.03.04.709535

**Authors:** Richard Mcnair, Carl A. Whitfield, Gowsihan Poologasundarampillai, Oliver E. Jensen, Igor L. Chernyavsky

**Affiliations:** Department of Mechanical, Materials and Manufacturing Engineering, University of Nottingham, University Park, Nottingham, NG7 2RD, UK; Department of Mathematics, University of Manchester, Oxford Road, Manchester, M13 9PL, UK; Faculty of Dentistry, Oral and Craniofacial Sciences, King’s College London, Great Maze Pond, London, SE1 1UL, UK; Maternal and Fetal Health Research Centre, University of Manchester, Oxford Road, Manchester, M13 9WL, UK

**Keywords:** human placenta, stereology, 3D imaging, mathematical modelling, diffusive transport

## Abstract

**Introduction:** Stereological estimates of villous membrane thickness and surface area are widely used to infer the diffusive exchange capacity of the human placenta. A key geometric determinant of exchange capacity can be expressed as an effective diffusive length scale. Here we combine virtual histological sections with computational modelling in realistic villous geometries to assess the accuracy of classical stereological estimates of this diffusive length scale.

**Methods:** Two terminal villi, reconstructed from three-dimensional imaging, were digitally sectioned to generate random two-dimensional geometries containing fetal capillaries and surrounding villous tissue. For each section, we simulated steady diffusive transport between the fetal-capillary and intervillous-space boundaries to obtain a physics-based diffusive length scale as a reference case. Using the same geometries, we applied standard line–intercept stereology to measure harmonic-mean barrier thickness and boundary–length densities, from which a stereological estimate of diffusive length scale was derived.

**Results:** Across both villi, stereology systematically overestimated the diffusive length scale by approximately 15−25%, depending on villus and section. We identified sources of this discrepancy, including interface curvature and assumptions underpinning the stereological correction factors, using idealised models of villous structure.

**Conclusion:** These findings highlight the need for stereological approaches that account for curvature when interpreting placental structure–function relationships.

## 1. Introduction

In the human placenta, maternal blood flows from spiral arteries into a cavity filled with a villous tree of vascularised fetal tissue. The flow around this tree transports nutrients, which diffuse across a thin tissue barrier into the fetal circulation, with waste products diffusing the other way (Baergen et al., 2022). The thickness of this barrier, and its spatial variation, are key determinants of the diffusive capacity of the placenta in health and disease (Barapatre et al., 2026; Sun et al., 2020; Bappoo et al., 2024).

Many important species, such as oxygen, passively diffuse across the barrier (Nye et al., 2018; Gill et al., 2011). This process can be modelled using standard diffusion equations involving diffusion coefficients of the various species in human tissue (Perazzolo et al., 2017). The diffusive capacity of a barrier is directly proportional to a diffusive length scale ℒ (Erlich et al., 2019; Jensen and Chernyavsky, 2019). The diffusive exchange capacity is defined as the total diffusive transfer rate *J*_*d*_ across the exchange barrier driven by a fixed concentration difference Δ*c*, which can be written in the form

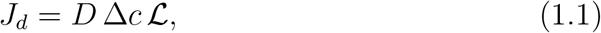

where *D* is the diffusion coefficient in tissue. Thus, ℒ summarises the morphometric component of diffusive exchange capacity, integrating both barrier thickness and its spatial variation. Historically (Mayhew et al., 1984, 2007), and in recent studies (Yong et al., 2023), ℒ has been approximated as the mean total exchange surface area divided by the harmonic mean of the barrier thickness, both of which can be estimated using stereological methods (Weibel and Knight, 1964).

Stereology is the technique for deriving information about higher dimensional structures from lower-dimensional data, for example two-dimensional sections of biological tissue. In the present context, placental tissue is typically fixed in formaldehyde and embedded in wax (see Table 2 for more details) before being sectioned to generate two-dimensional representations of the villous structure using stereological sampling designs (Jackson et al., 1985). To estimate barrier thickness, isotropic test lines, typically implemented using isotropic uniform random (IUR) sampling, are overlaid on the resulting sections, and the distribution of intercept lengths (chords) between the intervillous space and capillary interfaces is related to the harmonic-mean thickness via standard stereological formulae (Mayhew, 2006; Feneley and Burton, 1991), often implemented using dedicated stereological software. For calculating surface area or perimeter length, in three dimensions (3D) or two dimensions (2D) respectively, the intercept count of random lines per unit length of line can be related to the surface-area or perimeter density. These formulae are derived by calculating an expected value for the length of intersection/intercept count per unit length of a random line given the true thickness (Jensen et al., 1979; Gundersen et al., 1978). Many approximations are made in both the definition of the true thickness used in the derivation of the formulae, and in the calculation of this expected value (Mayhew, 2006). This traditional stereological approach was developed when only limited tissue sampling and imaging resolution were available, and it relies on geometric idealisations of the barrier that may not hold in the highly curved architecture of terminal villi. Today, high-resolution three-dimensional imaging and computational modelling (Chappell et al., 2023; Lewis and Pearson-Farr, 2020) allow us to evaluate how well classical stereological estimators recover diffusion-relevant quantities when applied to realistic villous geometries. This creates an opportunity to quantify possible systematic geometric bias, which is important when stereology-derived thicknesses and surface areas are propagated into estimates of placental diffusive capacity.

In this study, we use recent imaging data from micro-computed tomography (*μ*CT) and confocal laser-scanning microscopy (CLSM) derived datasets to sample randomised 2D geometries from villi by slicing the geometry into randomised sections and calculating ℒ_*p*_, the two-dimensional analogue of ℒ, to quantify diffusive exchange capacity on each section. Formal definitions of ℒ and ℒ_*p*_ are given in Supplementary Section S1. We compare these “physics-based” reference values to stereology-based estimates derived using established stereological approaches.

This study makes three contributions that directly inform interpretation of placental morphometric diffusion metrics. First, the physics-based values of ℒ_*p*_ provide new data to inform unbiased estimates of terminal villus diffusive capacity. Second, we quantify systematic error in stereological estimates and the uncertainty due to sampling. Third, we identify the geometric fea-tures of villous architecture, particularly curvature of thin exchange regions, that are a source of bias in stereological estimates of barrier thickness and diffusive length scale. Our aim is to evaluate how well classical stereological estimators recover the physics-based diffusive exchange capacity computed directly on the same geometries.

Throughout this work, we distinguish three different sources of error in stereological estimation of diffusive length scales or “exchange capacity”. The first is *stereology-formula* error, which is the bias introduced due to the assumption of an unbounded planar barrier made in the derivation of stereological formulae (as shown in Supplementary Section S2) linking harmonic-mean random intercept length with harmonic-mean barrier thickness. The second is *model-reduction* error, which is the discrepancy introduced when the diffusive length scale is approximated from the geometric proxy of an average perimeter or surface-area divided by a harmonic-mean barrier thickness. The third is *image-processing* error. The translation from raw images to labelled datasets inevitably involves errors and can produce significant artefacts, especially when quantifying surface areas and the thickness of thin barriers. Figure 1 illustrates these errors and where they arise in the pipeline from image to diffusive capacity estimate.

**Figure 1:**
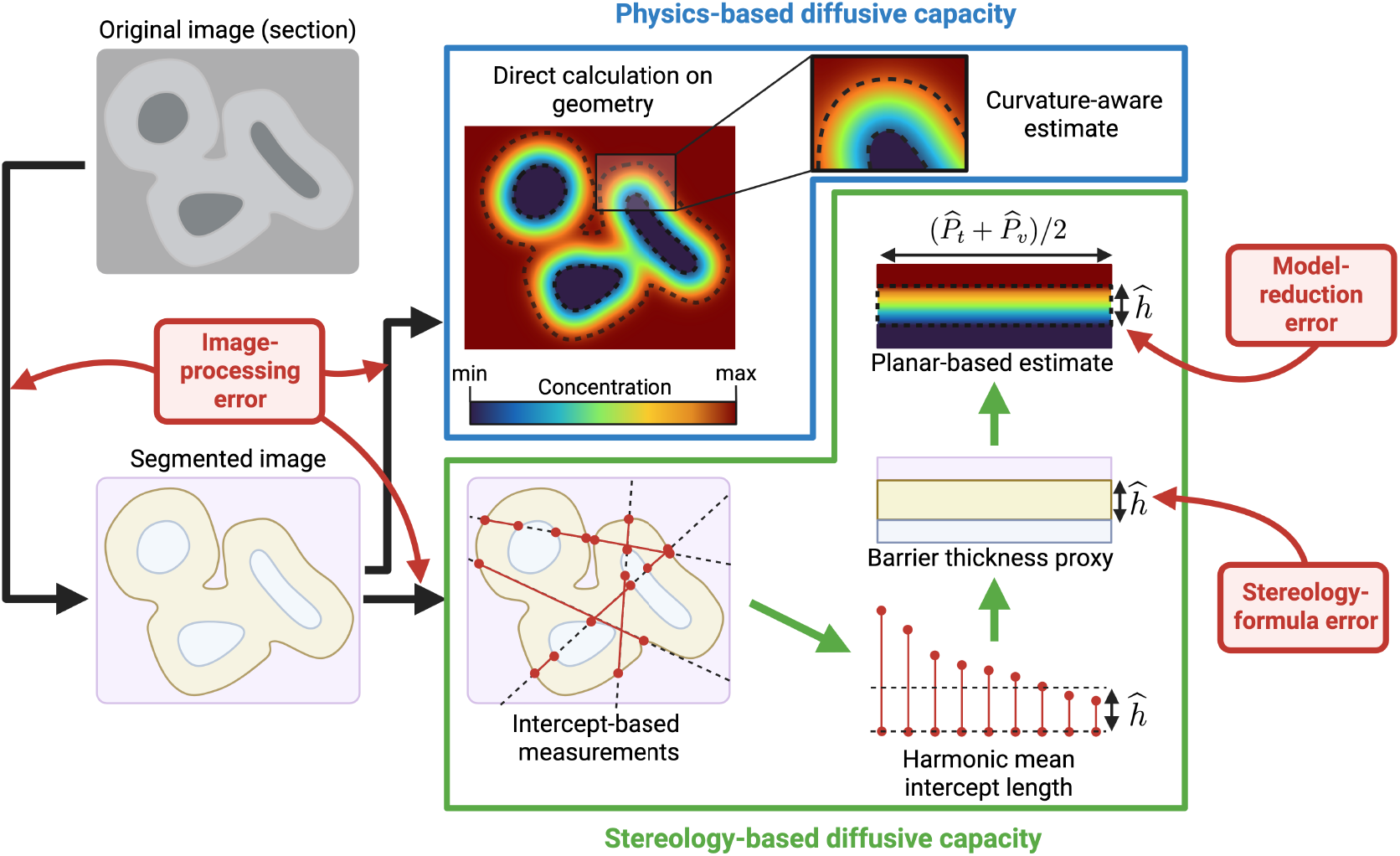
Schematic showing the two processes (physics- and stereology-based) to produce diffusive capacity estimates from a 2D section that we compare in this article. The red boxes illustrate the main sources of error that lead to disagreement between the two methods. “Image-processing error” includes errors due to image segmentation and the discretisation of geometries for both physics-based and stereology-based calculations. “Stereology-formula error” describes the use of formulae relating distribution of random intercepts to harmonic-mean thickness assuming small-gap planar surfaces, which may misrepresent irregular geometries (see equation S3.7). “Model-reduction error” describes the error introduced by calculating diffusive capacity from mean-surface area divided by harmonic-mean thickness assuming small-gap planar surfaces, as opposed to the physics-based method which calculates it directly from the geometry (see equation S3.8). Created in BioRender (https://biorender.com/uj4ic4a) with colour plots generated using VisualPDE (https://visualpde.com/).

The layout of this paper is as follows. In Section 2 we detail our method for directly calculating surface area and diffusive length scale from two terminal villus geometries. We then detail our methods for randomly sampling 2D sectioned geometries from the villi, and explain how we replicate stereological methods for calculating surface areas and harmonic mean thicknesses. We apply these methods to some simplified geometries where the thickness and diffusive capacity are known. In Section 3 we compare the stereologically-approximated surface areas and diffusive length scales with the physics-based values while comparing the results to experiments on simplified geometries. In Section 4, we discuss a possible source for the observed discrepancies in terms of surface curvature.

## 2. Model and methods

Table 1 defines all of the symbols used in the main text to represent measured quantities and model parameters. Table 2 provides a brief summary of the sample preparation and imaging protocols alongside the 3D morphological analysis of the two villous specimens (see Fig. 2).

**Table 1:** Symbols used in this study to represent various quantities, either defined as model outputs or measured as input parameters.

| Symbol | Description | Units |
| --- | --- | --- |
| Reported measures |  |  |
| $\mathcal{L}$ | Reference (physics-based) diffusive length scale of entire villus. | $\mu m$ |
| $\mathcal{L}_p$ | Reference (physics-based) diffusive length scale per unit depth of a sampled 2D section, defined in equation S1.3 of Supplementary Section S1. | — |
| $\hat{\mathcal{L}}_p$ | Stereologically-estimated diffusive length scale per unit depth of a sampled 2D section, defined in equation (2.4). | — |
| $P_t, P_v$ | Reference total perimeter of tissue (villus-IVS interface) and capillary (villus-capillary interface) respectively in a sampled 2D section. | $\mu m$ |
| $\hat{P}_t, \hat{P}_v$ | Stereologically-estimated total perimeter of tissue and capillary respectively, in a sampled 2D section. | $\mu m$ |
| $h$ | Reference (physics-based) effective barrier thickness in a 2D section, calculated as mean reference perimeter divided by $\mathcal{L}_p$. | $\mu m$ |
| $\hat{h}$ | Stereologically-estimated effective barrier thickness in a 2D section, calculated by equation (2.3). | $\mu m$ |
| Stereology parameters |  |  |
| $A_{ROI}$ | Area of region of interest (ROI) | $\mu m^2$ |
| $N$ | Number of chords (tissue intercepts) generated from the random lines | — |
| $I_t, I_v$ | Number of intercepts of generated lines with tissue and capillary surfaces respectively, for a given set of lines and 2D section. | — |
| $T_i$ | Length of the $i$th tissue chord (segment of a test line connecting IVS and capillary interfaces) | $\mu m$ |
| $L_T$ | Total length of line sections within $A_{ROI}$ | $\mu m$ |
| Idealised model parameters |  |  |
| $\kappa$ | Curvature of the circular interface used in Section 3.1. | $\mu m^{-1}$ |
| $d$ | Thickness of the planar-band and annular benchmark geometries used in Section 3.1. | $\mu m$ |

**Table 2:** Summary of imaging, sample preparation, and morphometric characteristics for the two terminal villus datasets analysed (Specimen 1: synchrotron *μ*CT; Specimen 2: confocal microscopy). Sources: ^a^ Tun et al. (2021); ^b^ Plitman Mayo et al. (2016); ^c^ Plitman Mayo et al. (2019), Erlich et al. (2019).

|  | Specimen 1 | Specimen 2 |
| --- | --- | --- |
| Modality | Synchrotron μCT | Confocal microscopy |
| Sample/Data ID | 123761 | 1 |
| Clinical characteristics | Term CS; healthy pregnancy | Term CS; healthy pregnancy |
| Fixation and embedding | Z7+PFA perfusion fixed; Batson's infiltrated via fetal side; critical point dried | Perfusion-fixed with formalin |
| Staining | — | FITC-conjugated Ulex lectin (perfused; vessels); DiI dye (immersed; syncytium) |
| Feature extraction software | Avizo | 3DSlicer; SolidWorks |
| Raw imaged volume ($\mu\text{m}^3$) | $\lesssim 10^{10}$ | $\sim 10^7$ |
| Voxel size: $x, y, z$ ($\mu\text{m}$) | 0.81, 0.81, 0.81 <sup>a</sup> | 0.28, 0.28, 2.00 <sup>b</sup> |
| Total specimen volume ($\mu\text{m}^3$) | $1.96 \times 10^5$ | $2.59 \times 10^6$ <sup>b</sup> |
| Capillary volume fraction | 0.34 | 0.28 <sup>b</sup> |
| Capillary-to-villous surface ratio | 1.01 | 1.05 <sup>b</sup> |
| Harmonic mean thickness ($\mu\text{m}$) | — | 8.37 <sup>c</sup> |
| Directly calculated diffusive length-scale $\mathcal{L}$ ($\mu\text{m}$) | $0.62 \times 10^4$ | $0.79 \times 10^4$ |

**Figure 2:**
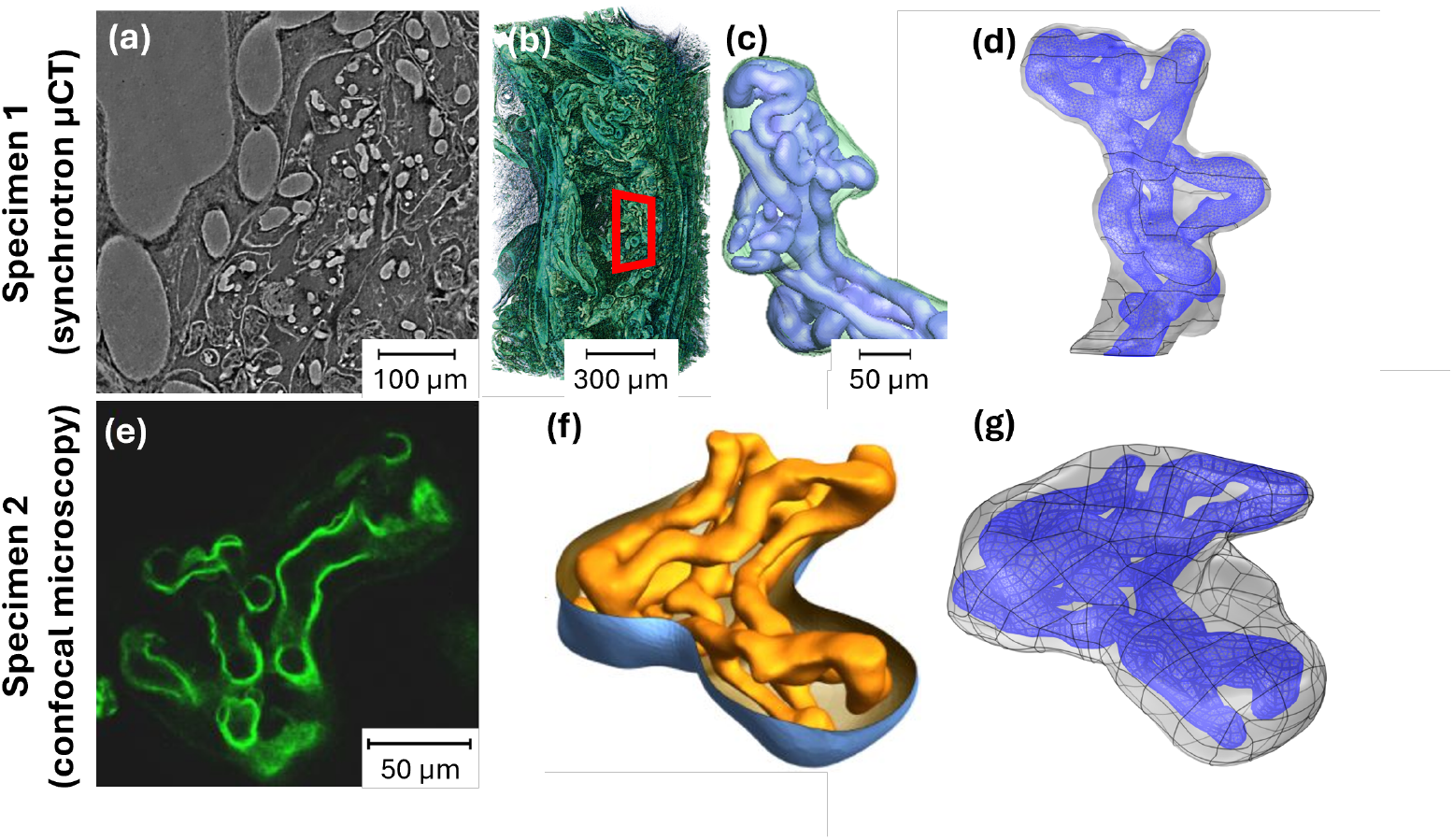
Creation of terminal villus datasets and computational meshes used for slicing and stereological analysis. (a–d) Villus 1 from synchrotron *μ*CT (adapted from Tun et al. (2021) under CC BY 4.0). (a) A representative 2D section showing villi and vessels. (b) 3D rendering of a smaller region of interest showing the complex placental architecture. (c) 3D rendering of a terminal villus and capillaries, taken from within the red box of (b). (d) Finite-element mesh of the terminal villus geometry imported into `COMSOL Multiphysics`. (e–g) Villus 2 from confocal microscopy (adapted from Erlich et al. (2019) under CC BY 4.0). (e) 2D confocal section of the villus. (f) Segmented surface geometry of the villus and capillary network. (g) Corresponding finite-element mesh in `COMSOL` used for random slicing and stereological analysis.

### 2.1. Approximation of stereological techniques on computational data

In this subsection, we describe how classical stereological measurements are reproduced on computationally-generated 2D sections of the villous geometries, enabling direct comparison with physics-based reference quantities.

We performed stereological experiments on two terminal villus geometries (Fig. 2), which we identify as villus 1 and villus 2. Each villus was imported into `COMSOL Multiphysics 6.3`^®^ (COMSOL AB, 2024) and enclosed within a rectangular bounding box (Fig. 3a,b) which is needed to calculate perimeters from stereologically obtained perimeter densities. Two-dimensional sections were generated by intersecting the villus and its bounding box with uniformly-randomly oriented planes, producing, for each villous specimen, a set of 20 2D-section geometries (Fig. 3c). Sections were retained only if the plane intersected both the tissue and capillary domains and for villus 2, sections intersecting a cropped edge were excluded. This yielded 15 valid sections for villus 1 and 8 for villus 2. Each section contains (i) the tissue boundary, (ii) the capillary boundary, and (iii) the bounding polygon that defines the region of interest (ROI). All implementation details required for reproducibility of the methods are provided in Supplementary Section S3. We perform the analysis here in two dimensions only, as it allows more data to be generated for comparison. The corresponding three-dimensional estimators are obtained by pooling intercept counts and intercept lengths over multiple sections, and differ only by standard numerical conversion factors (Gundersen et al., 1978). Thus, the stereology-formula and model-reduction type errors we find in two dimensions are expected to persist to the 3D case too.

**Figure 3:**
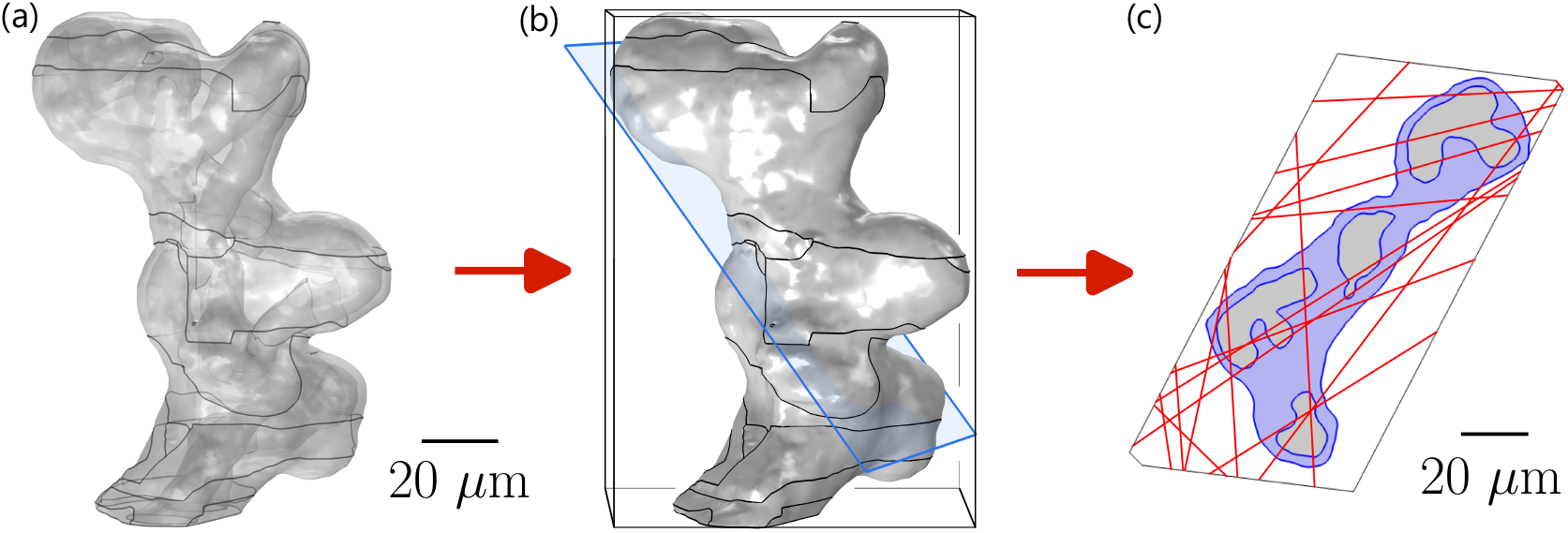
Overview of the stereological workflow. (a) Villus 1 geometry is imported into `COMSOL`. (b) A rectangular bounding box is drawn around the villus, and a random section is sampled through the villus and box. (c) The work plane yields a 2D section geometry consisting of the villus outline, capillary outline, and a bounding polygon defining the region of interest (ROI). Random test lines are overlaid on the section and used to estimate perimeter lengths and harmonic-mean barrier thickness.

For each section, COMSOL was used to compute reference values against which stereological estimates were compared. These included the total tissue and capillary perimeters, *P*_*t*_ and *P*_*v*_ respectively, measured directly from the section geometry, and the 2D diffusive length scale ℒ_*p*_ obtained by solving the steady diffusion equation on the same section. This diffusive length-scale is proportional to the diffusive capacity in two dimensions in the same way as ℒ in (1.1) is in three dimensions. This quantity is computed numerically (using finite-element methods in `COMSOL`) and is therefore subject to small discretisation errors; however, these are negligible relative to the geometric effects quantified in this study.

To present diffusion-based predictions in a form that is directly comparable to classical harmonic-mean thickness measurements, we define a reference ‘physics-averaged thickness’, *h*, for each two-dimensional section such that

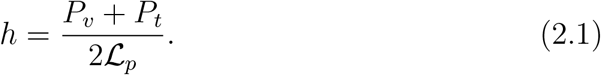

This is the thickness of a notional uniform planar barrier that would give the same overall diffusive exchange as the original section geometry under the same concentration difference.

For each section, a high-resolution image was exported together with its spatial dimensions. Each image was segmented into tissue, capillary, and intervillous space (IVS) within the ROI. Stereological estimators were then applied to each segmented section by overlaying random test lines on the ROI, as illustrated in Figure 3(c).

#### 2.1.1. Total perimeter and harmonic mean barrier thickness estimation from random lines

Perimeters were estimated from the number of boundary crossings made by 500 isotropic uniform random test lines overlaid onto each segmented section image. Let *I*_*t*_ and *I*_*v*_ denote the total number of intercepts between the test lines and the tissue–IVS boundary and the tissue–capillary boundary respectively. Let *L*_*T*_ denote the total length of all test-line segments lying within the ROI, and let *A*_ROI_ denote the ROI area in the section. The stereological estimates of the tissue and capillary perimeters on a section are (Romppanen and Collan, 1983; Cruz-Orive, 2017)

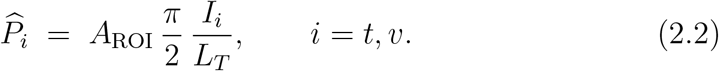

This is the standard isotropic uniform random line-intercept estimator for boundary length density in 2D. 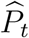 and 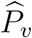 seek to estimate *P*_*t*_ and *P*_*v*_ respectively. We note that the derivation of (2.2) does not require assumptions of local planarity or small gap thickness and is valid for arbitrarily curved interfaces under isotropic uniform random sampling.

Barrier thickness was estimated from tissue chord lengths sampled along the random test lines. Each chord corresponds to a portion of a test line passing through tissue between IVS and capillary. The section-level stereological estimate of harmonic-mean barrier thickness is derived in Supplementary Section S2 as

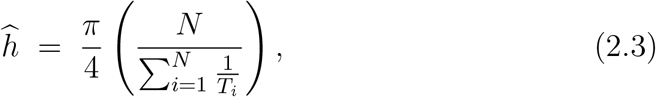

where *T*_*i*_ denotes length of the *i*th tissue intercept and *N* is the number of tissue intercepts generated from the random lines (the 3D relation can be found in standard references like Jensen et al. (1979)). In contrast to (2.2), geometrical assumptions of thin gap thickness and local planarity are made in the derivation of (2.3) as shown in Supplementary Section S2.

### 2.2. Stereological proxy for the diffusive length scale

In this subsection, we describe how the stereological estimates of perimeter and harmonic-mean thickness are combined to form the standard proxy for the diffusive length scale used in the literature. From the stereological measures above, we define a section-level estimate of the diffusive length scale per unit depth by combining perimeter and thickness estimates as

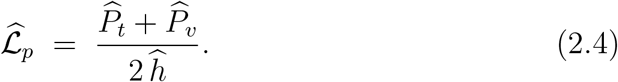

This proxy uses the arithmetic mean of the tissue and capillary perimeters, normalised by the harmonic-mean barrier thickness, which is the typical approximation used in literature (Mayhew et al., 1984, 2007; Yong et al., 2023). The resulting 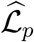values were compared directly against the reference ℒ_*p*_ values computed in `COMSOL` on the same sections.

Sampling uncertainty due to the finite number of random test lines was quantified using bootstrap resampling at the level of test lines within each section. For each section, repeated resampling with replacement produced empirical distributions for the estimated perimeters 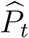, 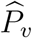, harmonic mean thickness 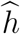, and diffusive length scale per unit depth 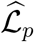, from which 95% confidence intervals were reported. Full details of the bootstrap procedure are given in Supplementary Section S3.

## 3. Results

### 3.1. Experiments on simplified geometries

To help interpret errors for predicting barrier thickness from stereological methods, we carried out two benchmark experiments using simplified synthetic images where the barrier thickness is known exactly. In both experiments, we generated labelled 2D images representing a tissue barrier separating a vessel region from the intervillous space, and then applied the random-line stereological thickness pipeline used in the main section experiments. Repeating the experiment many times allowed us to separate any bias from random variation caused by finite sampling. These experiments reveal errors of stereology-formula type only and thus help us understand the relative contribution of both stereology-formula and model-reduction errors in our experiments on the villus geometries.

First, we tested a planar barrier of constant thickness spanning the full width of the image (Fig. 4a). This geometry is designed to isolate the effect of finite barrier aspect ratio: when the barrier is not very thin compared with the overall field of view, stereological assumptions become less accurate. We repeated the experiment across a range of barrier thicknesses while adjusting the image resolution so that the barrier was always similarly well-resolved in pixels (Supplementary Section S3, Fig. S1). This allowed us to attribute any remaining error to geometry rather than pixellation.

**Figure 4:**
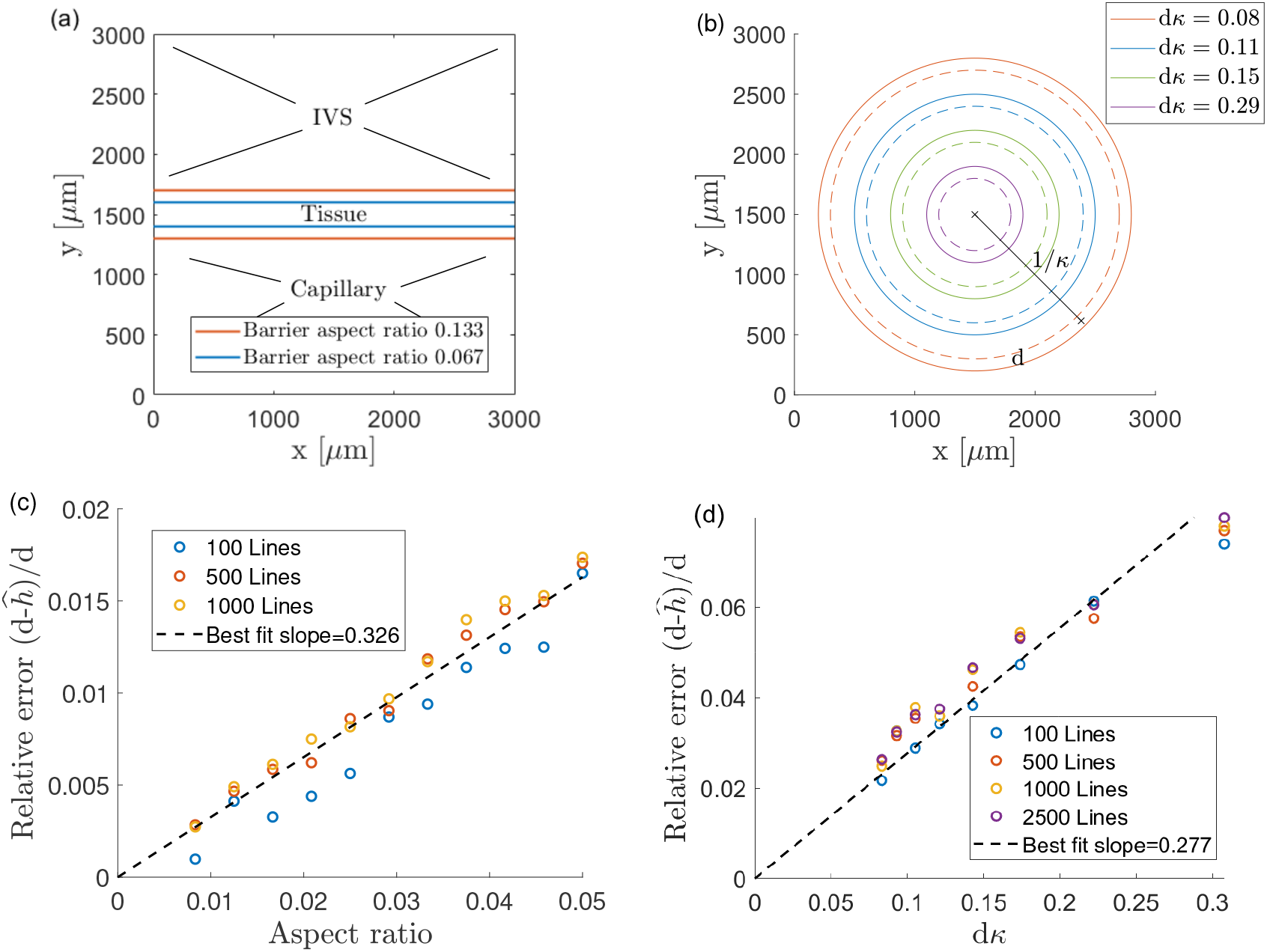
Experiments on simplified geometries. (a) Schematic of a finite-aspect-ratio barrier. Blue lines show a barrier of aspect ratio 0.067, red curves show a barrier with aspect ratio 0.133. (b) Schematic of the annular geometries considered. Solid curves denote the outer boundary and dashed curves the inner boundary. Colours indicate different barriers with radii of mid-circles 1*/κ* (equal to the inverse of curvature *κ*); the barrier thickness is a constant d across geometries. (c) Relative error in stereological approximations *h* of barrier thickness *d* versus barrier aspect ratio for the planar barrier geometries in experiments using 100, 500 and 1000 lines, showing convergence as the aspect ratio tends to zero. Best-fit slope shows approximate error as 0.326 times the aspect ratio. (d) Relative error in stereologically approximated thickness for the annular geometries against non-dimensional parameter d*κ*. A best-fit slope quantifies error as approximately 0.277d*κ*.

Second, we tested a curved barrier formed by two concentric circles, giving a tissue annulus of constant thickness but with varying curvature (Fig. 4b). By changing the circle radius while keeping the barrier thickness fixed, we generated barriers that ranged from weakly to strongly curved. This geometry isolates curvature-driven bias in harmonic-mean thickness estimation. As with the planar experiment, results were obtained by repeated random-line sampling, and the mean error was summarised across trials to identify how bias depends on curvature.

The model-reduction errors in circular and spherical geometries for ℒ_*p*_ and ℒ (reported in Supplementary Section S4) can be calculated analytically and are proportional to the dimensionless parameter (*dκ*)^2^, where *d* is the barrier thickness and *κ* is the curvature. In these experiments we isolate the stereology-formula errors and also study these in terms of *dκ* to directly compare their relative contributions.

Figure 4 reports the benchmark experiments on the simplified geometries. The planar experiment (Fig. 4c) demonstrates convergence of the stereological harmonic-mean thickness estimate as the barrier aspect ratio decreases, consistent with the thin-barrier assumptions underlying the stereology formulae. The concentric annulus experiment (Fig. 4d) shows an approximately linear increase in the relative thickness error with the dimensionless curvature parameter *dκ*, supporting the interpretation that curvature-driven deviations from the locally-flat idealisation systematically underestimate thickness and hence overestimate ℒ_*p*_. Across Figures 4 and 5, bootstrap confidence intervals quantify sampling variability arising from the finite number of random test lines, while the fitted slopes highlight systematic disagreement that persists beyond this sampling uncertainty.

**Figure 5:**
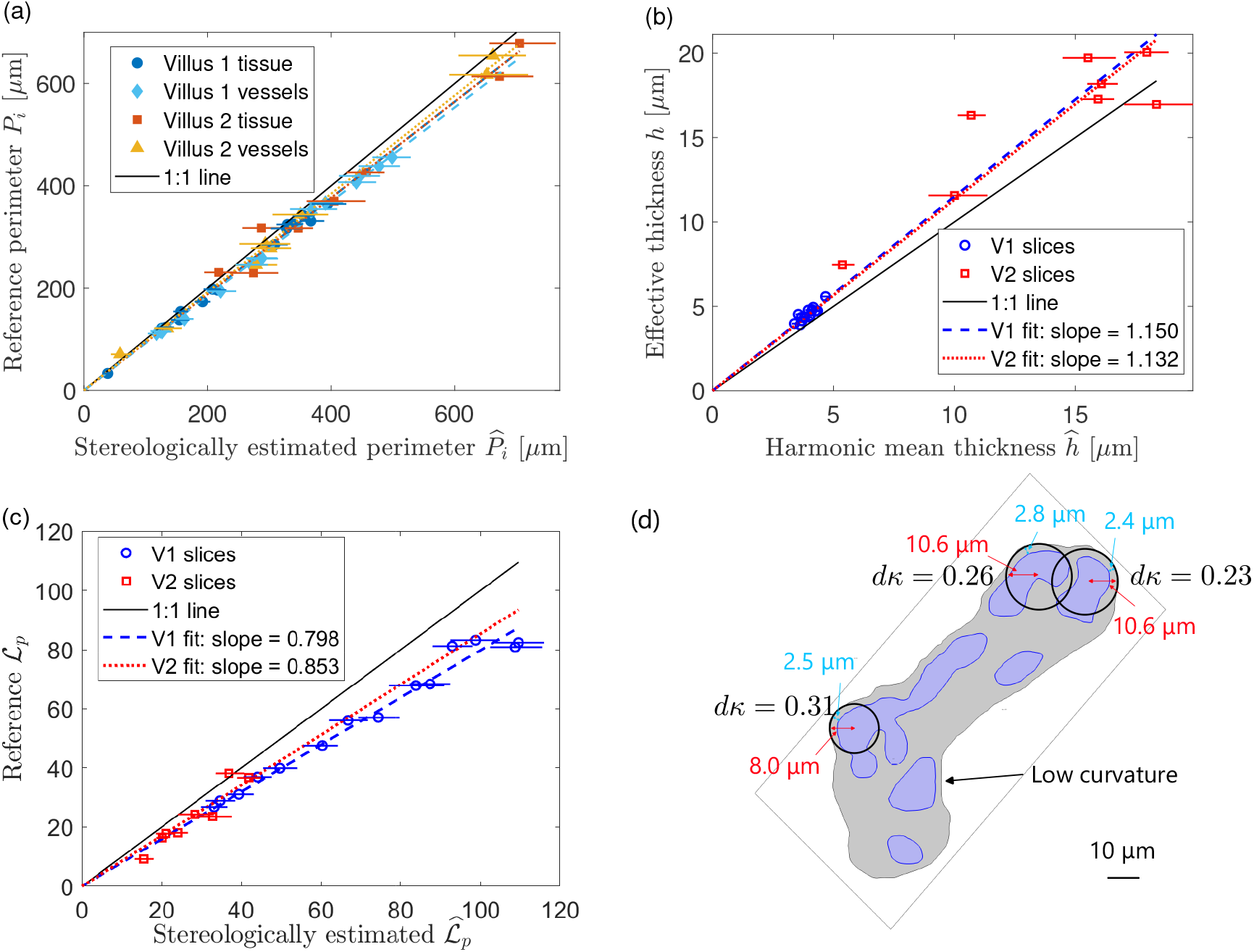
Results from the stereological experiments using images of villi. (a) Stereologically-estimated perimeter (*x*-axis) is plotted against the reference perimeter (*y*-axis). Best-fit dashed lines of slope 0.941, 0.922, 0.941 and 0.961 for villus 1 tissue, villus 1 vessel, villus 2 tissue and villus 2 vessel respectively compare the estimated and reference values. (b) The stereologically-estimated harmonic-mean thickness (*x*-axis) is compared with the physics-based thickness (*y*-axis). (c) The stereology-based estimate 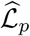 (*x*-axis), defined by equation (2.4) is compared with the reference diffusive length scale ℒ_*p*_ (*y*-axis). In (a)–(c), error bars show 95% bootstrap confidence intervals for the stereological estimates, and best-fit lines are included to highlight systematic disagreement. (d) A representative section with circles marking the thinnest regions of the tissue boundary, illustrating typical local curvatures.

These simplified geometries are designed to isolate specific sources of bias, such as finite aspect ratio and curvature, in a controlled setting where the true thickness is known exactly. It must be emphasised, however, that they do not capture the full morphological complexity of terminal villi, which exhibit tortuosity, branching, and spatially heterogeneous curvature.

### 3.2. Experiments on villus geometries

Figure 5 summarises the section-by-section stereological estimates compared with reference values computed directly from the `COMSOL` geometries. In Figure 5(a), stereological perimeter estimates show a small but consistent bias. Linear fits through the origin for the four data series (tissue and vessel perimeters in villi 1 and 2) have slopes 0.941, 0.922, 0.941 and 0.961, respectively, indicating that the computational procedure systematically biases the stereologically-estimated perimeters to be slightly larger than their reference values. A plausible source of this discrepancy is jaggedness of the tissue and vessel boundaries induced in the processing of the images, which increases the measured interface length relative to the underlying smooth geometry. Importantly, although the magnitude of this perimeter bias is similar to the other sources of bias described below, it is not in itself large enough to explain the bias in the calculation of ℒ_*p*_ which we describe next.

Figure 5(b) shows a systematic discrepancy in stereologically-estimated thickness compared with the “physics-based” effective thickness (that is, the uniform effective thickness that reproduces the same diffusion-derived exchange capacity as the original section geometry). The stereological harmonic-mean thickness is biased low compared with the physics-based estimates, with best-fit slopes of 1.150 for villus 1 and 1.132 for villus 2 (i.e. the reference thickness is *≈*10–15% larger than the stereological estimate across sections). This indicates that classical line-intercept stereology, when applied to these realistic villus sections, tends to underestimate the effective exchange-barrier thickness. Since the diffusive length-scale proxy 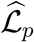 depends inversely on thickness, this systematic thickness bias dominates the overall behaviour of the stereological estimate 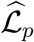. This means that stereological methods systematically underestimate the effective barrier thickness in these two realistic villous geometries, particularly in thin, highly curved regions that contribute most strongly to diffusive transport.

Figure 5(c) shows that stereology systematically overestimates ℒ_*p*_, with best-fit slopes (for physics-based versus stereological estimate) of 0.798 and 0.853 for villi 1 and 2, respectively. Equivalently, the stereological estimate is typically larger than the reference value by approximately 15–25%. Figure 5(d) highlights the highly curved, thin regions that contribute strongly to the harmonic-mean thickness. The thinnest regions in representative sections correspond to typical values of *dκ ≈*0.23–0.31, which contribute to the systematic overestimation of the diffusive length scale by stereological methods, and therefore the inferred diffusive exchange capacity, for these geometries. To summarise, using the idealised model-reduction error derived in Supplementary Section S4, a value of *dκ*=0.3 would give an error of *∼* 1–5%. Using Figure 4(d), the corresponding stereology-formula error is *∼* 10%.

## 4. Discussion

### 4.1. Principal findings

In this study, we used fully-segmented 3D villous geometries and numerical solutions of the diffusion equation to quantify the accuracy of classical stereological methods for estimating placental exchange barrier thickness and diffusive exchange capacity. By comparing stereological estimates obtained from virtual 2D sections with reference values calculated by `COMSOL Multiphysics`, we showed that conventional line-intercept–based stereology systematically overestimates the diffusive length-scale ℒ_*p*_ in 2D images. By extension, the overall villous diffusive length scale ℒ may also be overestimated. This bias was observed in two independent villous geometries obtained from distinct imaging modalities and segmentation pipelines, suggesting that it reflects generic geometric features of villous architecture rather than idiosyncrasies of a single specimen or modality.

Our analysis further indicates that this bias might be linked to local curvature of the fetal capillary–syncytiotrophoblast interface. The discrepancy we have observed in the calculation of the diffusive length scale arises from multiple sources. In particular, two curvature-linked effects are sufficient to generate significant bias even in the absence of image-processing errors. First, there is stereology-formula error. Even with exhaustive *in silico* sampling, the classical line-intercept correction factors are derived under idealised assumptions of locally planar, parallel interfaces, so they do not fully account for curvature-dependent terms. Second, there is model-reduction error. Approximating diffusive capacity using the ratio of mean surface area to harmonic-mean thickness is itself a geometric simplification, and becomes increasingly inaccurate when barrier thickness varies over a curved interface. Consequently, stereological estimates of ℒ derived from thickness and surface–area measurements may not converge to the numerically derived values even as sampling density increases. Together, these findings suggest that, while stereology provides useful relative comparisons between groups, its use to infer absolute diffusive capacity requires careful re-evaluation when curvature and thickness heterogeneity are substantial. We note, however, that geometric effects do not in general guarantee a bias of fixed sign. In more complex configurations of the exchange interface, it is conceivable that stereological estimates could either over- or under-estimate diffusive capacity. Nevertheless, across all cases examined in this study the dominant effect was a systematic overestimation. Establishing the generality of this behaviour across a wider range of geometries remains an important direction for future work.

### 4.2. Comparison with previous work

Stereological methods have been central to quantitative placental morphometry for several decades, underpinning much of our understanding of how villous structure supports feto-placental exchange. Numerous studies (e.g. Mayhew et al. 1984, 2007; Yong et al. 2023) have used point-counting and line-intercept techniques to estimate villous volume, capillary surface area, and thickness of the villous membrane, and then combined these measures into morphometric indices of diffusive capacity. In particular, the concept of a harmonic-mean barrier thickness, often estimated from line-intercept distributions, has been widely used as a surrogate for the effective path length for diffusion.

By explicitly solving the diffusion problem in realistic geometries, we show that the effective diffusive length depends not only on harmonic-mean barrier thickness and surface area, but also on how that thickness is distributed in space and curved in relation to the driving concentration gradient. Classical stereological formulae for the harmonic-mean thickness calculation effectively collapse these details into a planar approximation, which is violated in villi with tortuous capillaries, bulging syncytial knots, and irregular microvillus patterns.

The overestimation of ℒ_*p*_ that we observe is therefore consistent with the idea that stereological estimates of harmonic-mean thickness, derived from line intercepts across complex curved interfaces, do not fully capture the impact of geometry on diffusive pathways. Our curvature-based analysis suggests that systematic deviations from planarity accumulate into appreciable biases when propagated through the standard formulae linking thickness, surface area, and diffusive capacity.

### 4.3. Methodological implications for placental stereology

The most immediate implication of this work is methodological. Our results do *not* imply that stereology is invalid for studying placental structure, but they do highlight the following points.

1. Absolute values of diffusive exchange capacity derived from classical formulae should be interpreted with caution. In practical terms, the magnitude of bias we observe is potentially not negligible: stereology underestimates barrier thickness *h* by approximately 10–15% and overestimates ℒ_*p*_ by approximately 15–25% (Fig. 5b,c); this directly impacts absolute barrier-thickness values (Feneley and Burton, 1991; Ong and Burton, 1991) if the bias we observe in this study scales-up to whole organ measurements. Since morphometric formulations show diffusive capacity as proportional to the diffusion coefficient multiplied by an effective diffusive length scale, this implies that stereology-based capacity estimates could be inflated by a comparable percentage, even before considering additional experimental sources of error. This mat-ters because changes of similar magnitude in morphometric or diffusive-capacity measures are often interpreted as functionally significant in the placental literature (Mayhew et al., 1984, 2007; Feneley and Burton, 1991). More broadly, this suggests that absolute values of diffusive capacity reported in the literature should be interpreted with caution, as they may systematically overestimate true exchange capacity due to geometric bias. More work is needed to clarify these biases.
2. Relative comparisons are likely more robust than absolute values. Despite the bias, we found reasonably consistent relationships between stereological and numerical estimates across sections, indicating that stereology captures differences in barrier thickness and surface–area– to–volume ratios between regions or groups, provided sampling and analysis are identical.
3. The development of a method for correcting for curvature may improve stereological estimates. Our derivations show that curvature terms naturally enter the expressions linking geometric proxies to diffusive capacity (Supplementary Section S4) and curvature could be incorporated in stereological formulae for harmonic mean thickness. Although we have not yet distilled a simple, universally applicable correction factor, the analysis suggests a route towards modified stereological formulae that explicitly incorporate measures of interface curvature or shape.
4. The choice between stereological and 3D modelling approaches depends on the scientific objective. Stereology remains well suited to high-throughput analyses across large cohorts, where the primary aim is to compare relative differences between groups under consistent sampling protocols. In contrast, physics-based 3D modelling provides more direct access to structure–function relationships and is better suited to mechanistic studies or to establishing reference values for diffusive capacity. Our results suggest that these approaches should be complementary: 3D modelling can be used to quantify systematic geometric biases and inform the interpretation or calibration of stereological estimates, while stereology remains essential for population-scale estimates.

### 4.4. Limitations

Several limitations of our study should be acknowledged. First, our analysis is based on only two terminal villi, each reconstructed from a different imaging modality. While these villi are representative in terms of size and complexity, they cannot capture the full variability of villous architecture within and between placentas. Our conclusions about systematic bias should therefore be considered provisional until confirmed across a larger set of geometries spanning different gestational ages and clinical conditions.

Second, the notion of diffusive capacity used throughout this study is itself an idealisation. It is defined via a steady-state diffusion problem for a passive solute with a uniform diffusion coefficient, and neglects additional physiological processes such as binding, facilitated transport, and metabolic consumption. These assumptions are standard in morphometric analyses of placental transport and are inherent to the definition of diffusive capacity used in both stereological and modelling-based approaches.

Third, although we performed stereology on virtual sections to avoid cutting and processing artefacts, the image-processing and sampling protocols still introduce sources of error. Mis-classification at tissue boundaries, partial-volume effects at the voxel level, and choices about line density and step length can all affect estimated perimeter and thickness statistics. In experimental settings, additional biases due to fixation shrinkage, tissue deformation, and incomplete sampling of the villous tree are likely to be present.

Finally, our curvature analysis is currently restricted to relatively simple idealised geometries and local approximations. Extending this to more general surfaces, and developing practical curvature metrics that can be extracted from routine histological sections, remain important tasks for future work.

### 4.5. Future directions

Building on these findings, some avenues for future research are apparent. First, the stereology–simulation comparison should be extended to a larger cohort of villi and placentas, including both uncomplicated and pathological pregnancies. This would allow the magnitude and direction of stereological bias to be quantified across a wider spectrum of villous morphologies.

Second, it would be valuable to derive and validate simplified correction factors or lookup tables that map commonly measured stereological indices (such as harmonic-mean thickness and surface area) to effective diffusive exchange capacity for typical placental geometries. Such tools could make 3D, curvature-aware corrections accessible to groups without extensive modelling expertise.

### 4.6. Conclusions

In summary, by combining realistic villous geometries with computer simulations of the diffusive transport process, we have shown that classical stereological methods tend to overestimate the effective diffusive exchange capacity for solute transfer across the placental exchange barrier. This bias arises from assumptions of planarity and the neglect of curvature in the standard stereological formulae, and it persists even when sampling is dense and unbiased in two dimensions. While stereological methods are valuable for comparative morphometric studies, their use for deriving absolute diffusive exchange capacities should be treated with caution, ideally in conjunction with 3D modelling. Recognising and correcting for these geometric effects will improve the quantitative interpretation of placental structure–function relationships and may sharpen our understanding of how villous remodelling contributes to both normal and pathological feto-placental exchange.

## Acknowledgements

This work was partly supported by the Wellcome Leap *In Utero* programme. We thank Paul Brownbill, Angelos Evangelinos, Simon Cotter and Alexander Heazell (University of Manchester) and Michele Darrow and Avery Pennington (Rosalind Franklin Institute) for helpful discussions.

