## Supplementary material for "Physics-informed stereology for estimating placental diffusive exchange capacity"

---

#### S1. Definition of the diffusive length scale

The diffusive length scale for a barrier in three dimensions is defined by

$$\mathcal{L} = \int_S \mathbf{n} \cdot \boldsymbol{\nabla} c \; \mathrm{d}A, \tag{S1.1}$$

where $S$ is one of the surfaces enclosing the barrier, and $c$ satisfies

$$\nabla^2 c = 0 \quad \text{in } \Omega, \qquad c = 1 \text{ on } S_1 \quad \text{and} \quad c = 0 \text{ on } S_2. \tag{S1.2}$$

$\Omega$ is the region of space occupied by the barrier. In two dimensions, the diffusive length scale per unit length is defined as

$$\mathcal{L}_p = \int_P \mathbf{n} \cdot \boldsymbol{\nabla} c \;\, \mathrm{d}s \tag{S1.3}$$

where $P$ is one of the perimeters enclosing the barrier, d$s$ is an arc-length along the perimeter and $c$ satisfies Laplace's equation on the region in which $\Omega$ intersects the plane.

---

### S2. Relating sampled intercept lengths to the true mean thickness

This section derives the standard proportionality between the harmonic mean of sampled line intercept lengths and the perpendicular thickness of a membrane. The derivation assumes that the interfaces are locally parallel and planar. We present the 3D relation relevant to a membrane in space and the 2D relation relevant within a single planar section.

We assume that, over the spatial scale sampled by a probe line, the two bounding surfaces of the barrier are separated by a constant perpendicular distance $h$. Let $\Theta$ be the incidence angle between the probe direction and the normal of the planar interfaces such that $0 \leq \Theta \leq \pi/2$. Elementary geometry gives the relationship between the thickness $h$ and the measured intercept length $T$ along the probe as

$$\frac{1}{T} = \frac{\cos\Theta}{h}. \tag{S2.1}$$

This is where the assumption of parallel and planar surfaces/perimeters enters. The probability $g(\Theta)$ that a randomly oriented line will make an angle $\Theta$ with the interfaces is proportional to $\sin\Theta$ in three dimensions, but is uniformly weighted in two dimensions. However, the probability $p(\text{hit}|\Theta)$ that the line will make any intersection at all given $\Theta$, is proportional to $\cos\Theta$ in both cases.

Considering only lines which hit the interfaces, from Bayes's theorem the distribution $f$ of $\Theta$, given an intersection, is proportional to the product of these distributions

$$f(\Theta|\text{hit}) \propto g(\Theta)\, p(\text{hit}|\Theta). \tag{S2.2}$$

Therefore, the probability distribution for $\Theta$ is given by the normalised probability density functions

$$f_{2D}(\Theta|\text{hit}) = \cos\Theta \tag{S2.3}$$

for two dimensions, and

$$f_{3D}(\Theta|\text{hit}) = 2\cos\Theta\sin\Theta \tag{S2.4}$$

for three dimensions.

By substituting (S2.1) into (S2.3) and (S2.4) we obtain (showing 3D calculation outside of brackets and corresponding 2D calculation in brackets)

$$\begin{aligned}
\mathbb{E}\left[\frac{1}{T}\right] &= \int_0^{\pi/2} \frac{\cos\Theta}{h} f_{3D}(\Theta|\text{hit})\, d\Theta \quad \left(\int_0^{\pi/2} \frac{\cos\Theta}{h} f_{2D}(\Theta|\text{hit})\, d\Theta\right) \\
&= \frac{1}{h}\int_0^{\pi/2} 2\cos^2\Theta \sin\Theta\, d\Theta \quad \left(\frac{1}{h}\int_0^{\pi/2} \cos^2\Theta\, d\Theta\right) \\
&= \frac{2}{3h} \quad \left(\frac{\pi}{4h}\right),
\end{aligned} \tag{S2.5}$$

and thus

$$h = \frac{2}{3}T_h \quad \left(\frac{\pi}{4}T_h\right). \tag{S2.6}$$

where $T_h$ is the harmonic mean of random intercept lengths. This is an exact relation under the assumption of parallel, planar surfaces/perimeters but is only an approximation otherwise (Gundersen et al., 1978).

### S3. Methodology for simulated stereological experiments

This section describes the full 2D workflow used to estimate, via stereology methods, the tissue–IVS and tissue–capillary perimeters in each section and the harmonic-mean exchange-barrier thickness, together with bootstrap confidence intervals. These steps introduce sources of image-processing error, as defined in the main text, which may affect both perimeter and thickness estimates.

#### *S3.1. Method for obtaining randomly sectioned geometries*

In COMSOL, a rectangular bounding box is drawn around each villus. This is necessary as stereological methods for calculating surface area find surface area density and so a fixed volume must be imposed to calculate surface areas. Random 2D-sectioned geometries are sampled by a well-known method (Muller, 1959). First, three independent standard normal variables are sampled

$$[\hat{\mathbf{e}}_x, \hat{\mathbf{e}}_y, \hat{\mathbf{e}}_z] \sim [\mathcal{N}(0,1), \mathcal{N}(0,1), \mathcal{N}(0,1)], \tag{S3.1}$$

where $\mathcal{N}(0,1)$ denotes the standard normal distribution. Then, a unit normal vector is formed by normalising this sample,

$$\mathbf{n} = \frac{[\hat{\mathbf{e}}_x, \hat{\mathbf{e}}_y, \hat{\mathbf{e}}_z]}{||[\hat{\mathbf{e}}_x, \hat{\mathbf{e}}_y, \hat{\mathbf{e}}_z]||}. \tag{S3.2}$$

Next, a uniformly-random plane orthogonal to $\mathbf{n}$ is sampled from the set of all such planes that intersect the bounding box.

The resulting 2D geometries contain up to three families of curves: the curves that define the bounding box projected onto the plane; the curves which define the tissue surface; and the curves which define the vessel surfaces. In COMSOL a calculation of $\mathcal{L}_p$ was made on each section by first solving the 2D Laplace equation between the tissue and vessel perimeters and then integrating the result along the tissue perimeter according to (S1.3). The total perimeter length of the tissue $P_t$ and vessels $P_v$ were calculated using COMSOL's measurement tools.

*S3.2. Input data and physical scaling*

The sectioned 2D images were exported as $3000 \times 1500$ pixel jpegs, along with a measurement which defines the physical length represented by each side of each square pixel. The segmentation algorithm produces a label image $LAB$ with four classes:

- 0: outside the region of interest (ROI)
- 1: tissue
- 2: capillary
- 3: intervillous space (IVS)

Only pixels with labels in $\{1, 2, 3\}$ are inside the ROI.

*S3.3. Segmentation algorithm*

The algorithm consists of two stages: detection of the edges of the ROI and classification of pixels within the ROI into tissue, capillary, and IVS.

*Detection of the ROI polygon (projection of inner box to the section plane)*

1. The RGB image is converted to grayscale.
2. The black features in the image (the axes and the bounding box) are identified using an automatic thresholding step that picks out dark pixels.
3. After identifying non-dark pixels, MATLAB's `imclearborder` function is applied, which deletes any connected non-dark region that reaches the outer edge of the image. The remaining non-dark region is therefore the enclosed interior of the ROI, surrounded by the dark frame.

4. The polygonal border of the ROI is detected using MATLAB's `bwboundaries` function and stored as a closed polygon with $K$ vertices by listing the pixel coordinates $\{(x_k, y_k)\}_{k=1}^{K}$ around its edge.
5. The ROI area is computed directly from the label image as

$$A_{\mathrm{ROI}} = N_{\mathrm{ROI}} L^2, \tag{S3.3}$$

where $N_{\mathrm{ROI}}$ is the number of pixels within the ROI polygon.

*Classification within the ROI into tissue, capillary, and IVS*

1. The RGB image is lightly denoised using MATLAB's bilateral filter (`imbilatfilt`), applied separately to each colour channel to reduce pixel-level noise while preserving sharp boundaries.
2. The image is transformed into CIELab colour space, providing the lightness channel $L^*$ and chromaticity channels $a^*$ (green–red) and $b^*$ (blue–yellow).
3. To distinguish IVS (bright regions) from tissue and vessels (darker regions), we compute an automatic threshold on the $L^*$ values within the ROI using MATLAB's `graythresh` function. Pixels with $L^*$ below this threshold are labelled as non-white.
4. Vessels are identified inside this object mask by $k$-means clustering in the $(a^*, b^*)$ plane. The vessel cluster is selected as the one with lower mean $b^*$ (more blue).
5. Tissue is defined as the object mask excluding capillaries, and IVS is defined as ROI pixels not assigned to tissue or capillary.
6. Labels are assigned as IVS $\to 3$, tissue $\to 1$, capillary $\to 2$, and outside ROI $\to 0$.
7. Overlays of the segmented boundary on top of the original images are inspected for quality. A small amount of post-processing is applied by eroding the capillary mask by 1-2 pixels to visually give a clearer fit.
8. A final smoothing step is applied to reduce jagged label boundaries and reduce image-processing error as much as possible.

*S3.4. Random-line generation*

Random test lines are generated so that their orientations are isotropic and their placements are uniform with respect to translation over the ROI. For each random line

1. Sample an orientation angle

$$\theta \sim \mathrm{Unif}[0, \pi).$$

Define the unit normal and unit direction vectors

$$\mathbf{n}(\theta) = (\cos\theta,\ \sin\theta), \qquad \mathbf{u}(\theta) = (-\sin\theta,\ \cos\theta).$$

2. To determine the range of line positions that intersect the ROI, each polygon vertex is projected onto $\mathbf{n}(\theta)$ and the minimum and maximum values across all vertices are taken

$$s_{\min}(\theta) = \min_{k=1,\dots,K} \big(\mathbf{n}(\theta)\cdot(x_k, y_k)\big), \qquad s_{\max}(\theta) = \max_{k=1,\dots,K} \big(\mathbf{n}(\theta)\cdot(x_k, y_k)\big).$$

3. Sample an offset (signed distance) uniformly from this interval:

$$s \sim \mathrm{Unif}\big[s_{\min}(\theta),\ s_{\max}(\theta)\big].$$

4. The infinite line is defined implicitly by

$$\mathbf{n}(\theta) \cdot (x, y) = s.$$

5. Intersect this infinite line with the ROI polygon to yield a line segment within the ROI.

This procedure produces isotropic orientations and a uniform distribution of line offsets for each $\theta$, which is the standard construction for IUR lines within a bounded domain.

*S3.5. Test system: isotropic uniform random lines and discretisation*

For each section, $N_{\text{lines}}$ isotropic uniform random (IUR) test lines are generated, and the portion of the line lying within the ROI is traversed at a fixed step size of 0.25 pixels. This yields:

- the in-ROI test length for each line in $\mu$m,
- a one-dimensional sequence of labels along the line at uniform arclength increments,
- label transitions used to count interface intercepts,
- tissue chord lengths used for thickness estimation.

The total test-line length in the section is

$$L_T = \sum_{j=1}^{N_{\text{lines}}} \ell_j. \tag{S3.4}$$

*S3.6. Perimeter estimation from intercept counts*

Two interfaces are considered:

- tissue–IVS interface (labels 1 and 3), with intercept count $I_{31}$,
- tissue–capillary interface (labels 1 and 2), with intercept count $I_{12}$.

An intercept is counted whenever adjacent samples along a test line cross the relevant interface (both directions are counted equally). Per-section estimates of tissue–IVS perimeter and tissue–vessel perimeter respectively are obtained by multiplying by the ROI area

$$\widehat{P}_t \;=\; A_{\mathrm{ROI}} \frac{\pi}{2} \frac{I_{31}}{L_T}, \qquad \widehat{P}_v \;=\; A_{\mathrm{ROI}} \frac{\pi}{2} \frac{I_{12}}{L_T}. \tag{S3.5}$$

*S3.7. Thickness estimation from lineal intercepts*

Thickness is estimated from tissue chords extracted along the sampled IUR lines. Each tissue chord is defined as a contiguous run of label 1 (tissue) along a line that is flanked by IVS and capillary (in either order) along that line, corresponding to a local sequence 3–1–2 or 2–1–3. The physical chord lengths $\{T_i\}$ are computed from the number of steps in the run, the step size (0.25 pixels), and the pixel scale $L$.

The harmonic mean of the chord lengths is

$$H(\{T_i\}) \;=\; \left( \frac{1}{N} \sum_{i=1}^{N} \frac{1}{T_i} \right)^{-1}. \tag{S3.6}$$

where $N$ is the number of intercepts generated. The section-level harmonic-mean thickness estimate is then

$$\widehat{h} \;=\; \frac{\pi}{4} H(\{T_i\}). \tag{S3.7}$$

*S3.8. Section-level estimate of the diffusive length proxy*

For each section we compute diffusive length-scale per unit depth

$$\widehat{\mathcal{L}}_p \;=\; \frac{\widehat{P}_t + \widehat{P}_v}{2\,\widehat{h}}, \tag{S3.8}$$

and physics-based thickness

$$h \;=\; \frac{P_t + P_v}{2\,\mathcal{L}_p}. \tag{S3.9}$$

*S3.9. Bootstrap confidence intervals by resampling lines*

Sampling uncertainty arising from the finite number of random test lines is quantified using a nonparametric bootstrap performed at the level of test lines for each section.

For each section, the implementation stores per-line quantities:

- $I_{31,j}$: tissue–IVS intercepts contributed by line $j$,
- $I_{12,j}$: tissue–capillary intercepts contributed by line $j$,
- $\ell_j$: in-ROI test length for line $j$,
- $\{t_k\}_j$: the set of tissue chord lengths contributed by line $j$.

A bootstrap replicate is formed by resampling the $N_{\text{lines}}$ lines with replacement. For replicate $b$, totals are computed as

$$I_{31}^{(b)} = \sum_{j\in\mathcal{S}_b} I_{31,j}, \quad I_{12}^{(b)} = \sum_{j\in\mathcal{S}_b} I_{12,j}, \quad L_T^{(b)} = \sum_{j\in\mathcal{S}_b} \ell_j, \tag{S3.10}$$

and the perimeter estimates $\widehat{P}_t^{(b)}$, $\widehat{P}_v^{(b)}$ are obtained using the formulae above. The thickness estimate $\widehat{h}^{(b)}$ is obtained by pooling the chord lengths contributed by the resampled lines and applying the estimator above. The length proxy is then computed as

$$\widehat{\mathcal{L}}_p^{(b)} = \frac{\widehat{P}_t^{(b)} + \widehat{P}_v^{(b)}}{2\,\widehat{h}^{(b)}}. \tag{S3.11}$$

Confidence intervals are obtained from the empirical bootstrap distributions. A nominal 95% confidence interval is taken as the 2.5th and 97.5th percentiles.

*S3.10. Benchmark experiments on simplified geometries*

To help interpret the stereological errors observed on realistic villus sections, we also performed two benchmark experiments on simplified synthetic geometries. In each case we construct a labelled 2D image with a known barrier thickness, apply the same stereological thickness estimator as used in the main experiments, and compute their relative error. These simplified tests allow us to isolate two expected sources of bias: finite aspect ratio, and curvature.

*Finite-aspect-ratio barrier (planar band)*

We first consider a horizontal planar barrier of constant thickness spanning the full image width. This geometry isolates the effect of barrier aspect ratio. A square field of view of physical side-length

$$L_{\text{phys}} = 3000\ \mu\text{m} \tag{S3.12}$$

is discretised into a $M \times M$ pixel image. The pixel size is therefore

$$\Delta x = L_{\text{phys}}/M. \tag{S3.13}$$

For each trial and each tested thickness, we construct a horizontal band centered in the image that represents a tissue barrier between IVS and vessel regions. The pixels are labelled following the scheme in S3.3.

A range of thicknesses $d$ of the barrier are tested, and the corresponding aspect ratio is defined as

$$\text{A}_R = d/L_{\text{phys}}. \tag{S3.14}$$

To limit any error due to pixellation effects, the image resolution $M$ is adjusted with thickness so that the barrier remains similarly well-resolved for different aspect ratios

$$M \approx M_0 \frac{d_{\max}}{d}, \tag{S3.15}$$

where $M_0$ is a fixed resolution parameter and $d_{\max}$ is the largest tested thickness.

For each synthetic image, the stereological harmonic-mean thickness estimator is applied using $N_{\text{lines}}$ random lines as described in S3.7. For each barrier thickness, the relative error reported between the exact thickness and the estimated thickness in Figure 4(c) of the main paper is

$$\varepsilon_{\text{A}_R} = \frac{d - \widehat{h}}{d}. \tag{S3.16}$$

The experiment is repeated over many independent trials in order to estimate the mean bias as a function of aspect ratio and to visualise the scatter due to random-line placement.

*Curvature bias (concentric circular interfaces)*

To isolate curvature effects, we next consider a barrier between two concentric circles with fixed thickness but varying radius. In this case the barrier is locally curved, with curvature characterised using the mid-circle radius.

A square field of view of the same physical side-length $L_{\text{phys}} = 3000\ \mu\text{m}$ is discretised. For each geometry we specify an outer diameter $D_{\text{outer}}$ and an inner diameter

$$D_{\text{inner}} = D_{\text{outer}} - 200\ \mu\text{m}, \tag{S3.17}$$

so that the actual barrier thickness is constant across all cases:

$$d = \frac{D_{\text{outer}} - D_{\text{inner}}}{2} = 100\ \mu\text{m}. \tag{S3.18}$$

We test several outer diameters so that the curvature increases as the outer diameter decreases. This setup represents IVS outside both circles, tissue in the annular region and vessel inside the smaller circle, and the pixels are once again labelled according to the scheme from S3.3.

For each annulus, the same stereological harmonic-mean thickness estimate $\widehat{h}$ is computed using random lines as in S3.7. To express curvature effects in a dimensionless form, results are plotted against $d\kappa$, where $\kappa$ is the curvature of the circular interface. Since $\kappa = 1/R_d$ for a mid-circle of radius $R_d = (D_{\text{inner}} + D_{\text{outer}})/4$, we compute

$$d\kappa = \frac{d}{R_d}. \tag{S3.19}$$

The reported relative error is

$$\varepsilon_{d\kappa} = \frac{d - \widehat{h}}{d}. \tag{S3.20}$$

We repeat the experiment over many independent trials and compute the mean error as a function of $d\kappa$. A best-fit line constrained to pass through the origin is also plotted to summarise the approximately linear dependence of error on $d\kappa$ (Fig. 4(d) of the main paper).

These two benchmark experiments provide controlled reference cases for interpreting thickness-estimation bias: the planar band isolates finite-aspect-ratio effects, while the annulus isolates curvature-driven deviations from the locally-flat assumption implicit in the stereological correction factors.

*S3.11. Quantification of error due to pixellation*

The benchmark experiments in S3.10 are performed on synthetic label images and therefore introduce an additional numerical error associated with discretising a smooth geometry on a pixel grid which contributes to image-processing error. To quantify the magnitude of this pixellation error, we considered a single planar-band barrier of fixed aspect ratio 0.0667, and repeated the stereological thickness estimation while systematically increasing the image resolution.

For a series of resolutions (number of pixels across the square domain), we generated 100 independent realisations of the random-line sampling procedure using 200 isotropic uniform random test lines per trial. For each trial we computed the relative thickness error

$$\frac{d - \widehat{h}}{d}, \tag{S3.21}$$

where $d$ is the true barrier thickness and $\widehat{h}$ is the stereological harmonic-mean thickness estimate (Eq. S3.7). Figure S1 summarises the distribution of the relative error at each resolution using boxplots, demonstrating convergence of the estimator as number of pixels increases. This analysis confirms that, above a sufficiently fine resolution, the remaining bias in the planar-band experiment is not sensitive to pixellation.

### S4. Relationship between $\mathcal{L}$, surface area divided by thickness and curvature (analytical derivation of model-reduction error on concentric circles and spheres)

For an annular region in 3D (2D) between two concentric spheres (circles) with radii $R_1 > R_2$, the solution to Laplace's equation for dependent variable $c$ in the annular region according to $c = 1$ on the outer surface (perimeter) and $c = 0$ on the inner surface (perimeter) is given by the well-known solution (Crank, 1979)

$$c = \frac{R_1(r - R_2)}{r(R_1 - R_2)} \qquad \left(c = \frac{\ln(r/R_2)}{\ln R_1/R_2}\right) \tag{S4.1}$$

for $R_2 < r < R_1$. Taking the radial component of the gradient of $c$ and integrating it around the outer surface (perimeter) gives

$$\mathcal{L} = \frac{4\pi R_1 R_2}{R_1 - R_2} \qquad \left(\mathcal{L}_p = \frac{2\pi}{\ln R_1/R_2}\right). \tag{S4.2}$$

Calling $d = R_1 - R_2$, and assuming $d \ll R_1$ so that $\ln(R_2/R_1) = -d/R_1 - d^2/2R_1^2 + \ldots$, and also defining the mean radius as $R_d = (R_1 + R_2)/2$, (S4.2)

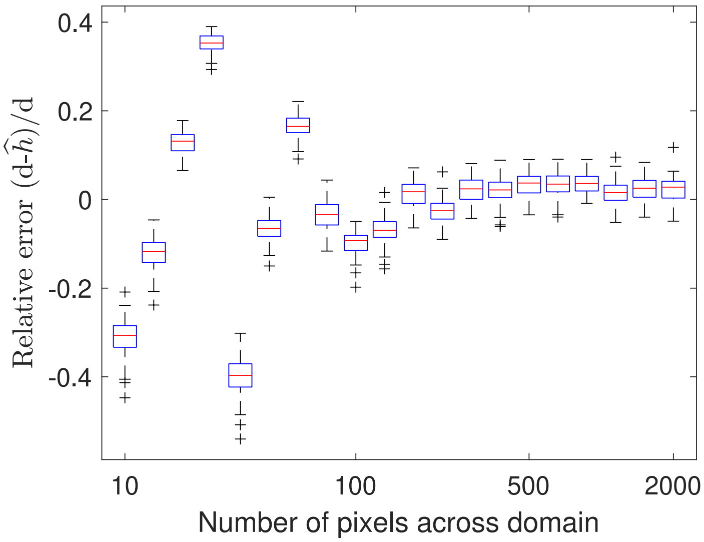


Figure S1: Convergence of stereological thickness estimation with image resolution for a planar-band barrier of fixed aspect ratio 0.0667. For each number of pixels across the square domain, the harmonic-mean thickness estimator was applied using 200 isotropic uniform random test lines, repeated over 100 independent trials. Boxplots show the distribution of relative error $(d - \widehat{h})/d$ at each resolution converging towards a constant.

is approximated as

$$\mathcal{L} = \frac{4\pi R_d^2}{d}\left(1 - \frac{(d\kappa)^2}{4}\right) \qquad \left(\mathcal{L}_p = \frac{2\pi R_d}{d}\left(1 - \frac{(d\kappa)^2}{12}\right)\right), \tag{S4.3}$$

where $\kappa = 1/R_d$. In the 2D case, the mean perimeter is $(P_1+P_2)/2 = (2\pi R_1 + 2\pi R_2)/2 = 2\pi R_d$; however in the 3D case, the mean surface area depends on curvature and is $(S_1+S_2)/2 = (4\pi R_1^2 + 4\pi R_2^2)/2 = 4\pi R_d^2(1+(\kappa d)^2/4)$. Then (S4.3) becomes

$$\mathcal{L} = \frac{(S_1+S_2)}{2d}\left(1 - \frac{(d\kappa)^2}{2}\right) \qquad \left(\mathcal{L}_p = \frac{(P_1+P_2)}{2d}\left(1 - \frac{(d\kappa)^2}{12}\right)\right), \tag{S4.4}$$

showing that the prefactors (mean surface area or mean perimeter divided by thickness) overestimate $\mathcal{L}$ or $\mathcal{L}_p$ irrespective of the sign of the curvature, in these simplified geometries.
